## Supplementary material for "Genome-Scale Modeling of *Rothia mucilaginosa* Reveals Insights into Metabolic Capabilities and Therapeutic Strategies for Cystic Fibrosis": Figure_S2.pdf

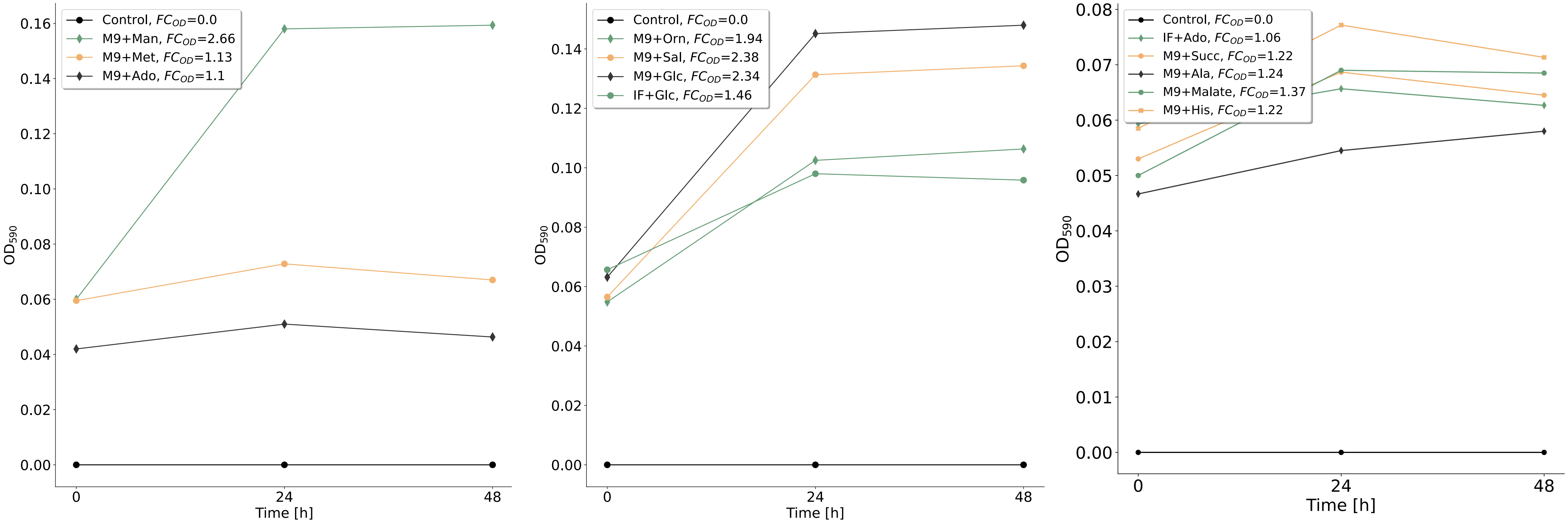


| | **M9+Man** | **M9+Met** | **M9+Ado** | **M9+Orn** | **M9+Sal** | **M9+Glc** | **IF+Glc** |
|---|---|---|---|---|---|---|---|
| BIOLOG | G | NG | NG | G | G | G | G |
| Stat. test | *** | ns | ns | * | ** | *** | * |

| | **IF+Ado** | **M9+Succ** | **M9+Ala** | **M9+Malate** | **M9+His** |
|---|---|---|---|---|---|
| BIOLOG | NG | NG | NG | NG | NG |
| Stat. test | ns | ns | ns | ns | ns |
