## Supplementary figures and images for "Genome-Scale Modeling of *Rothia mucilaginosa* Reveals Insights into Metabolic Capabilities and Therapeutic Strategies for Cystic Fibrosis"

### Figure_S1.pdf

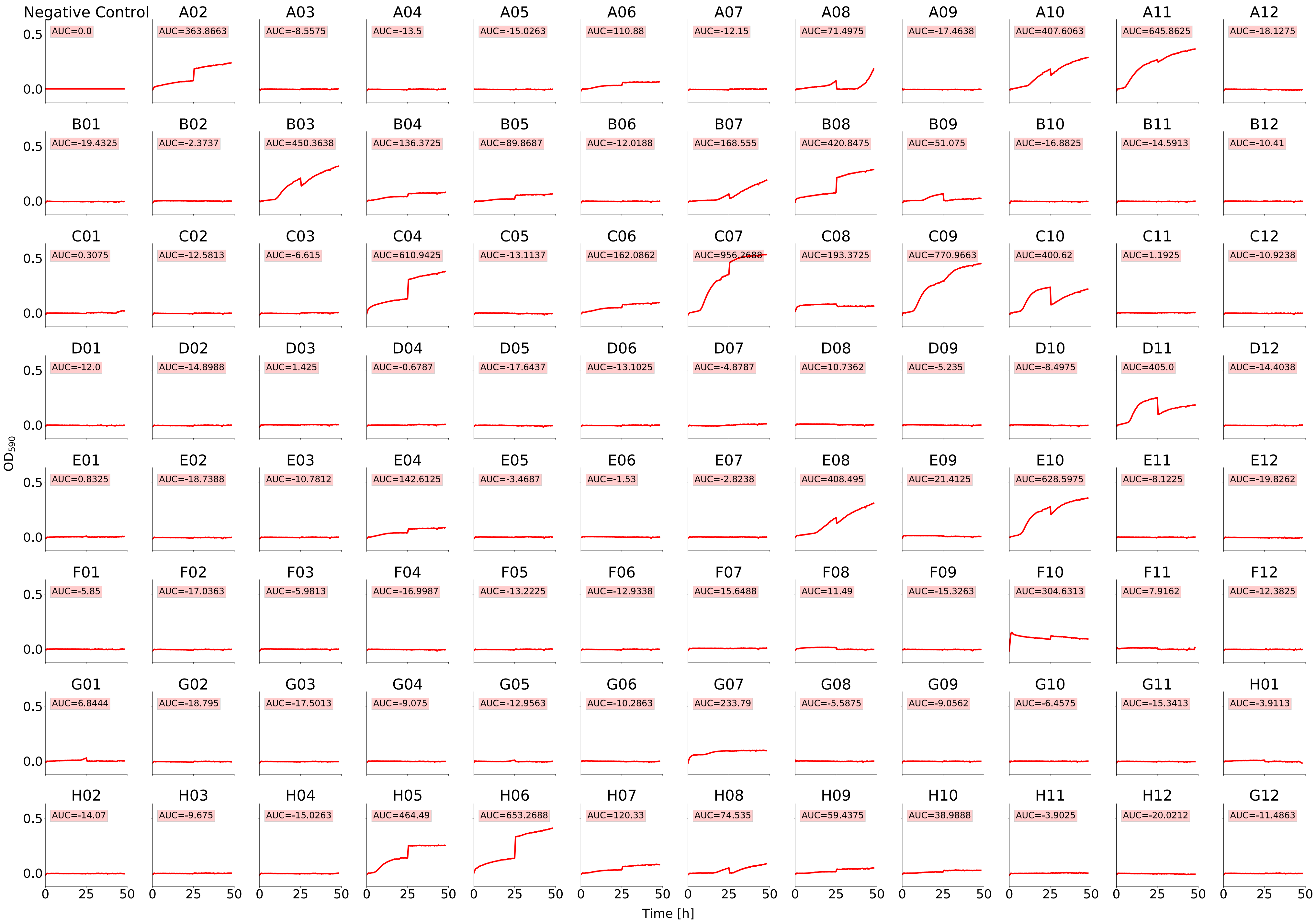

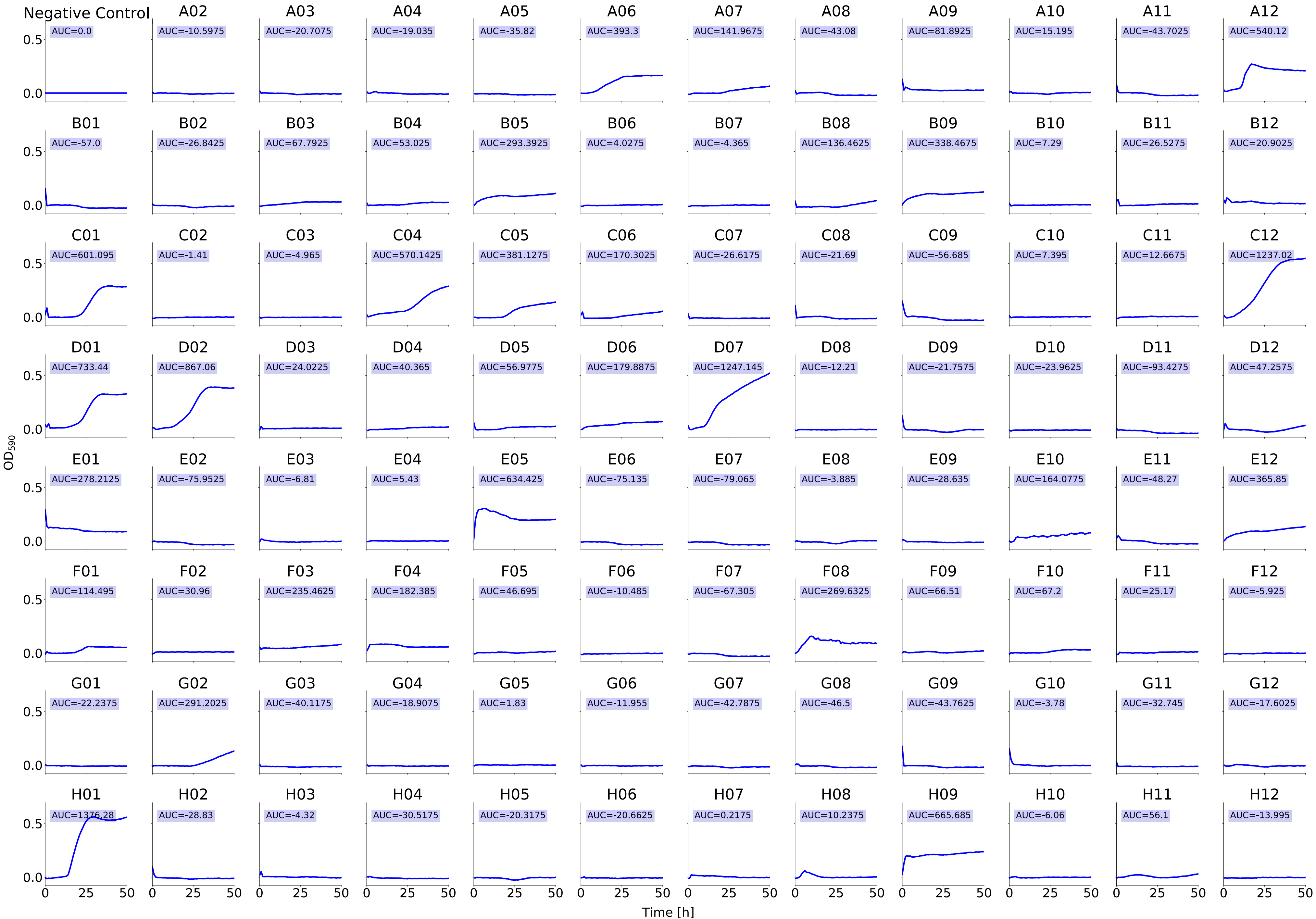

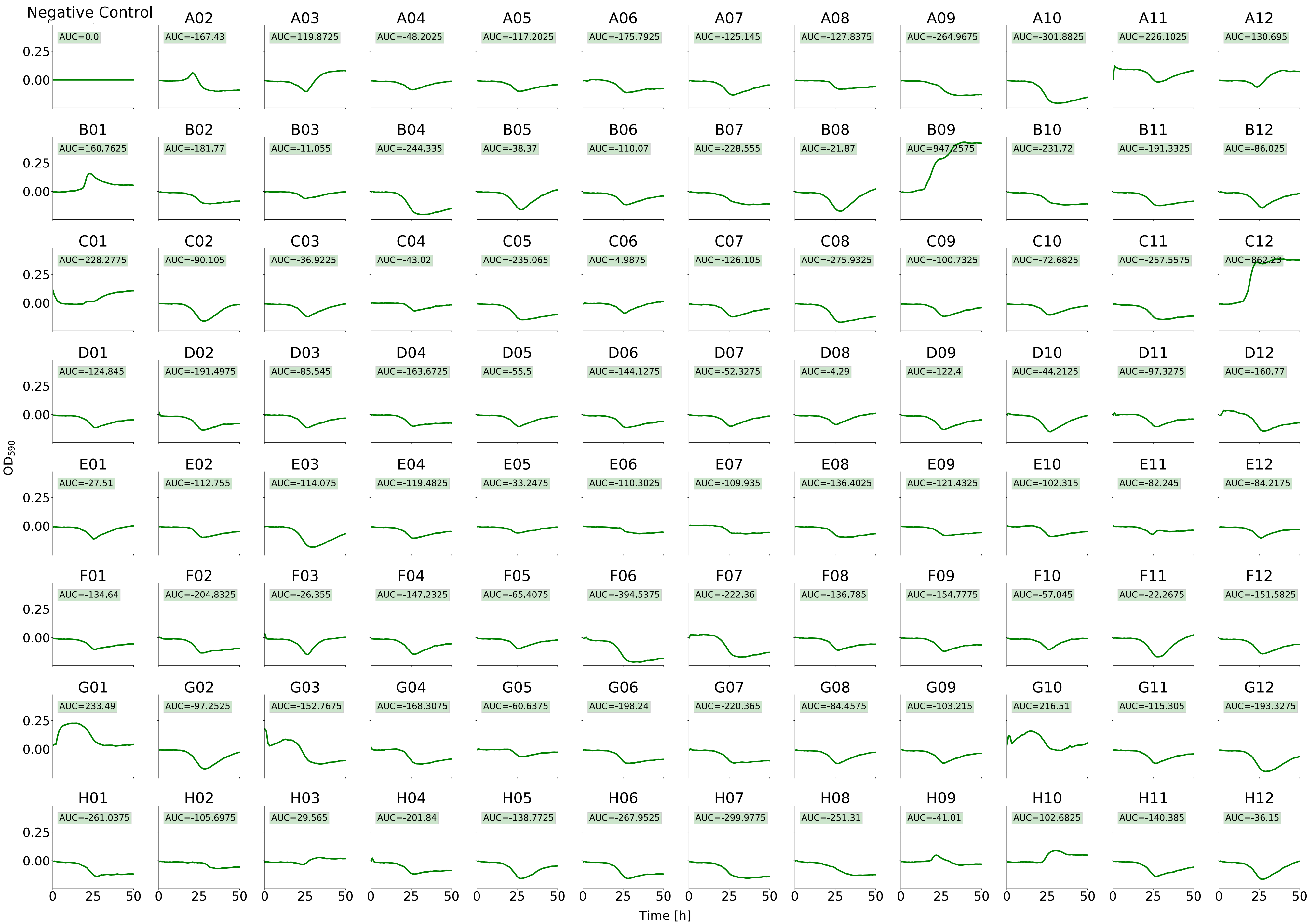

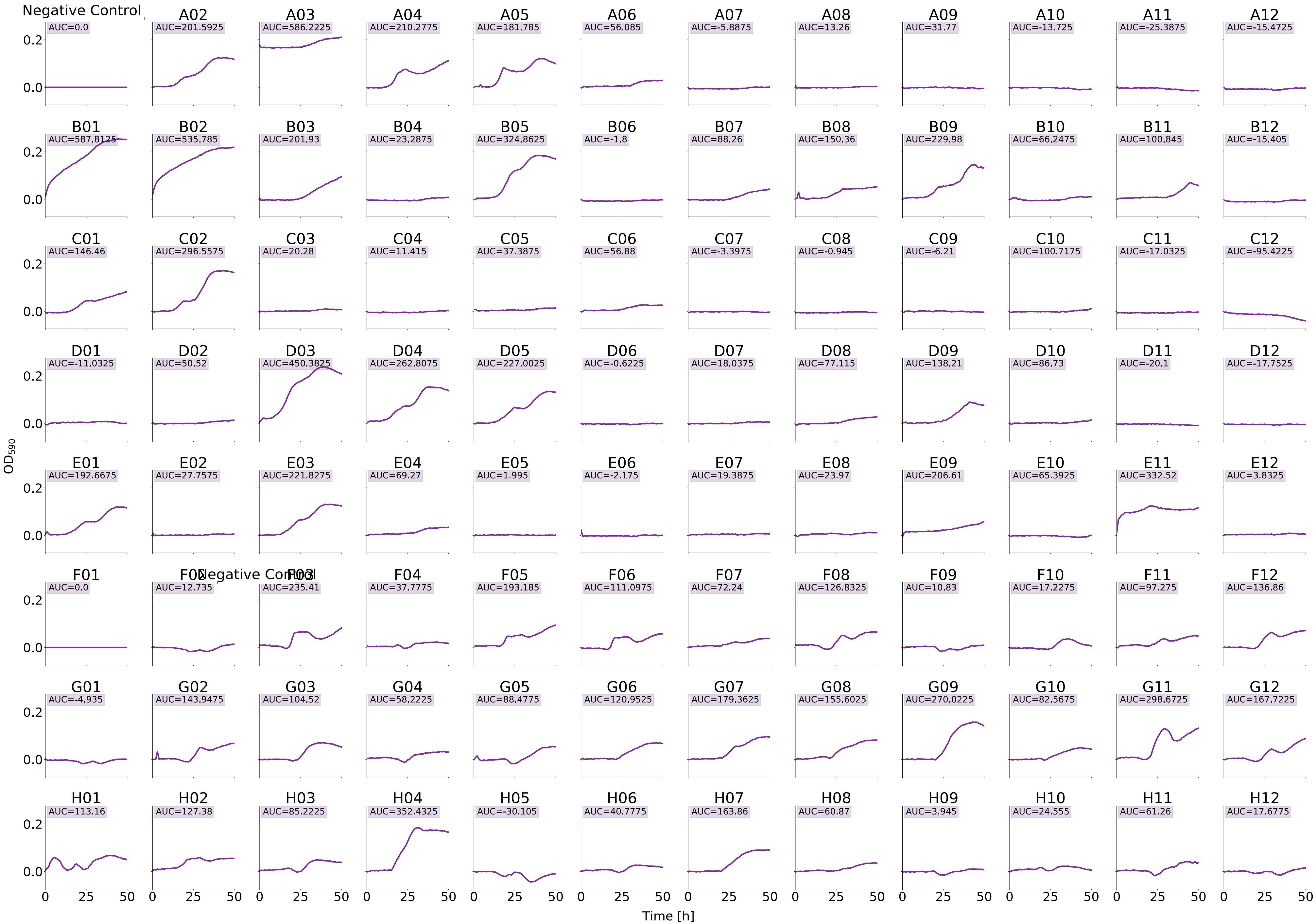

### Figure_S3.pdf

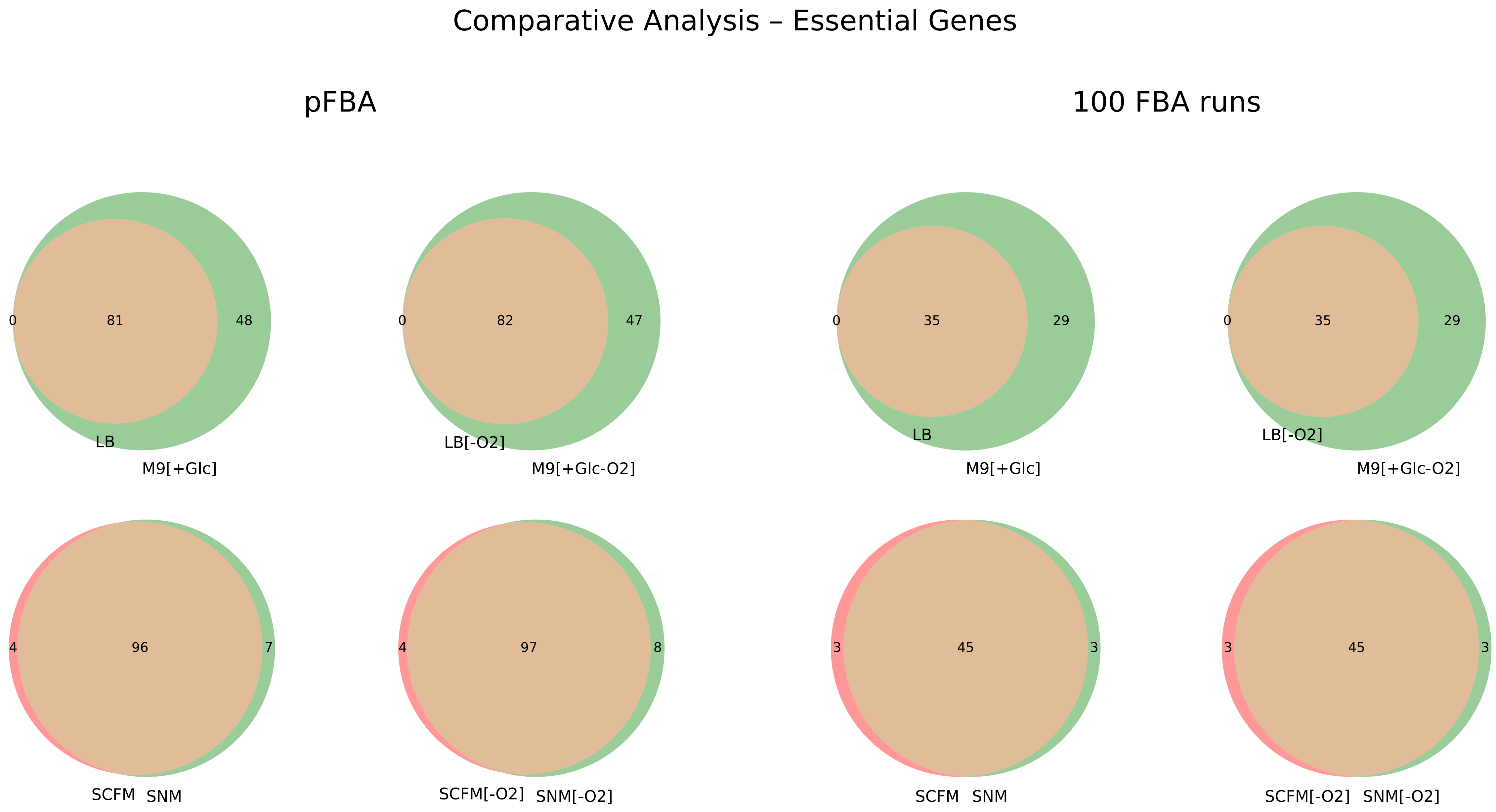

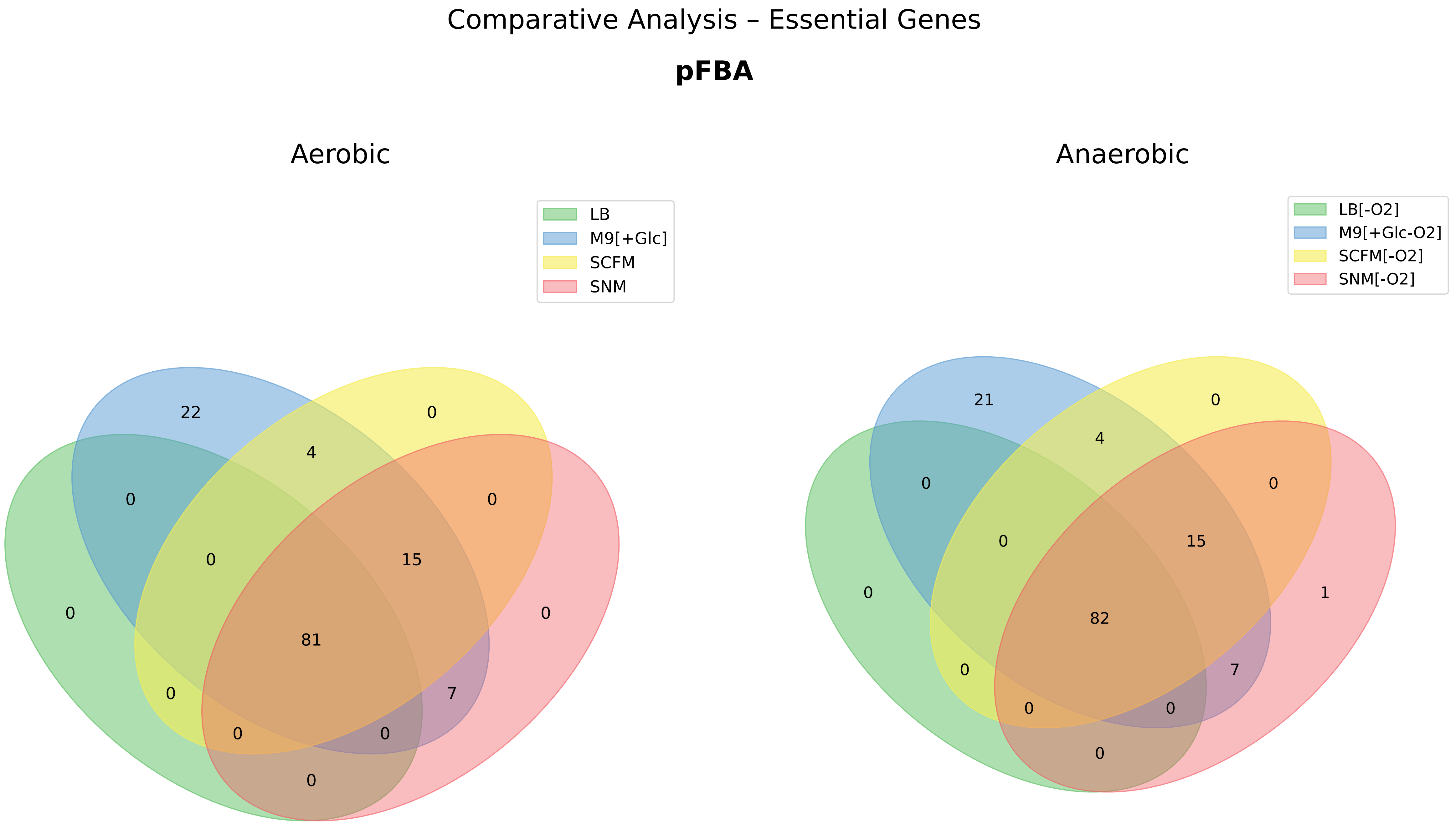

Comparative Analysis – Essential Genes

**100 FBA runs**

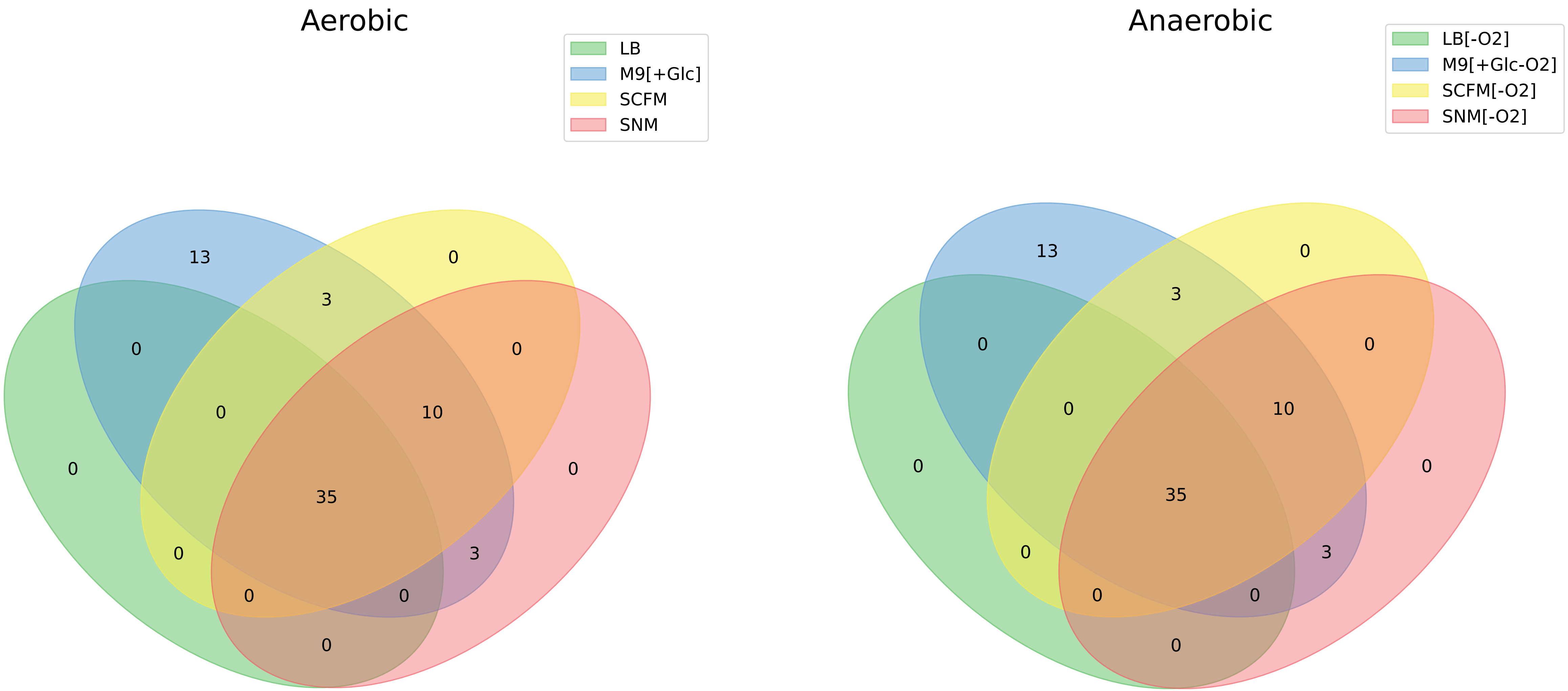

### Figure_S4.pdf

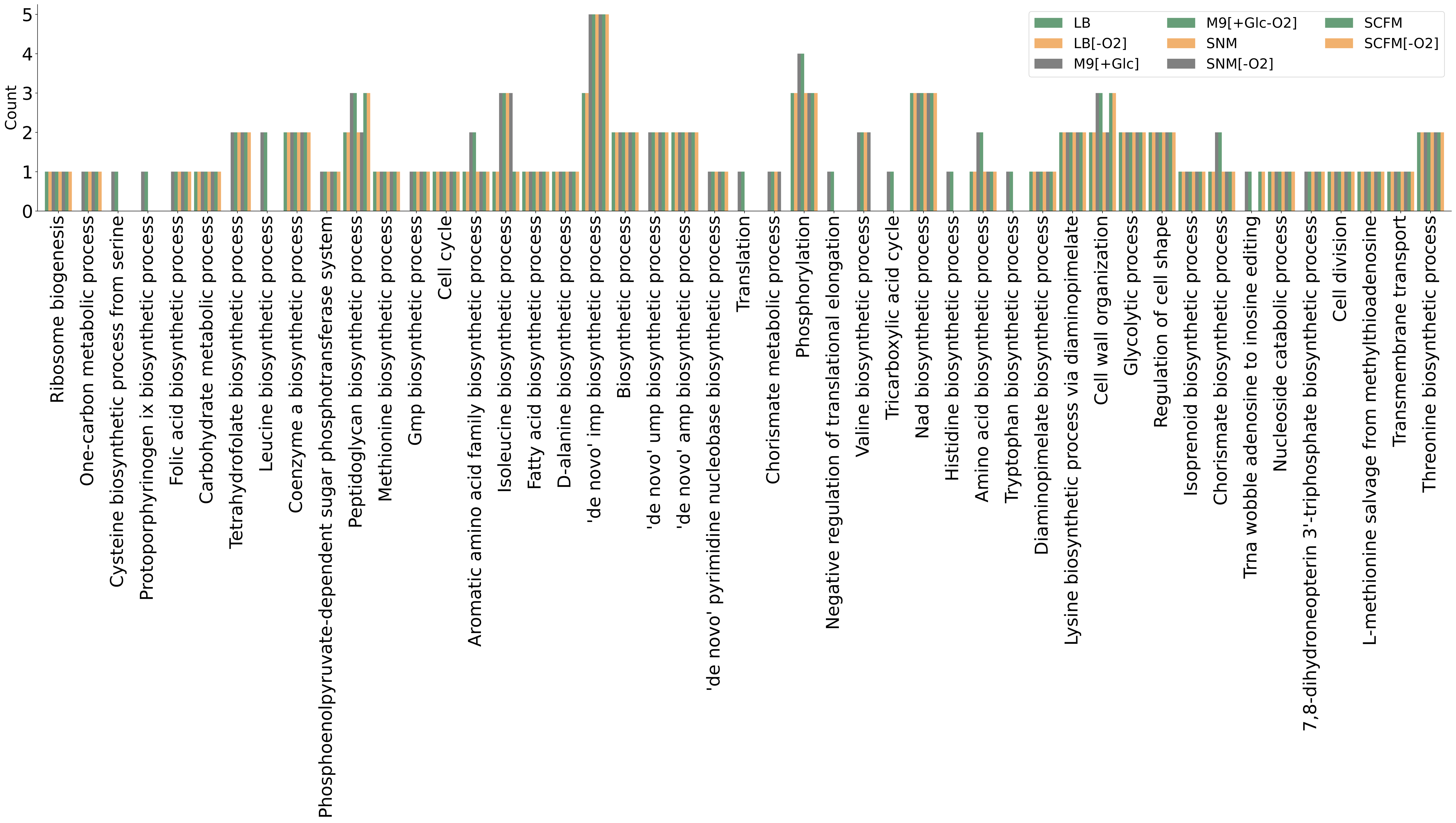
